# Expert-guided multi-objective optimization: an efficient strategy for parameter estimation of biological systems with limited data

**DOI:** 10.1101/2025.06.03.656243

**Authors:** Léa Da Costa Fernandes, David Bernard, François Pérès, Paul Monsarrat, Béatrice Cousin, Sylvain Cussat-Blanc

## Abstract

Calibrating biological models is challenging due to high-dimensional parameter spaces and the limited availability of reliable experimental data. In this study, we propose a hybrid calibration framework that integrates expert knowledge into a multi-objective optimization process using NSGA-II algorithm. Our approach combines hard constraints derived from biological measurements with soft constraints encoding qualitative domain expertise, such as expected curve shapes or event timing. This dual-constraint strategy guides the search toward biologically plausible parameter sets while preserving flexibility and interpretability. We demonstrate the effectiveness of our method on a benchmark model of skin wound healing, comparing it to standard and unconstrained optimization strategies. Results show that incorporating expert guidance significantly improves the biological relevance of simulated dynamics and mitigates overfitting, especially in underdetermined or uncertain settings. The framework is flexible, iterative, and generalizable, offering a principled way to leverage domain knowledge for model calibration in complex biological systems.

## Introduction

Computational or *in-silico* modelling in life sciences employs a diverse array of strategies to capture the complexity of biological systems across multiple scales. They can be broadly categorized into three main families of strategies [1]. First, mechanistic or determinist models use mathematical equations-such as ordinary or partial differential equations-to represent and predict the dynamic behavior of biological systems based on known biological mechanisms [1]. Second, data-driven models leverage statistical, machine learning, or other artificial intelligence techniques to uncover patterns and make predictions from large-scale biological data, such as genomics or proteomics datasets [2]. Third, hybrid and multi-scale models combine mechanistic and data-driven methods, or integrate different types of models (e.g., agent-based, Boolean, or constraint-based), to capture biological phenomena across multiple organizational levels-from molecules to whole organisms [1, 2]. This enables a more comprehensive and realistic simulation of biological systems and their emergent properties. Together, these general modelling strategies provide complementary perspectives for understanding, predicting, and manipulating complex life science phenomena.

Calibration is a critical step in the development of mechanistic and hybrid models, as it involves estimating the values of unknown parameters to ensure the model accurately reproduces biological data. The robustness and predictive accuracy of such models depend strongly on the quality of this calibration process [3]. While some parameter values can be extracted from existing literature, many remain unknown due to biological model specificity, experimental limitations or biological variability, some of which are not biologically accessible. Moreover, even when literature-derived values are available, they often depend on specific experimental conditions or biological contexts that may not align with the system under study. As a result, direct parameter estimation from biological data becomes necessary. This task is particularly challenging due to the high dimensionality of parameter spaces and the nonlinear relationships between parameters and model outputs [4, 5]. A poorly identified range of parameters can make calibration step more challenging.

Addressing this issue also requires the integration of additional sources of information beyond raw data. Expert knowledge, often qualitative in nature, can provide critical guidance in parameter estimation. For example, expert input can define expected behaviors, plausible parameter ranges, or curve profiles for variables. Numerous computational methods have been developed to address parameter estimation challenges, leveraging a range of optimization algorithms. For relatively simple biological models, classical curve-fitting techniques based on linear or nonlinear regression provide an effective solution [6]. More advanced approaches include gradient descent methods [7, 8], which iteratively refine parameters to minimize error, and metaheuristic techniques such as simulated annealing [9, 10, 11], which are particularly useful for navigating complex search spaces. Bayesian optimization [12], employing tools like Gaussian processes or machine learning emulator functions [13], excels in scenarios where model evaluations are computationally expensive. Other probabilistic methods, such as Markov Chain Monte Carlo [14, 15], allow for robust exploration of parameter distributions. Artificial neural networks have also been applied to parameter estimation tasks, leveraging their ability to approximate complex relationships between variables [16]. However, these algorithms typically require sufficient biological data to compare model simulations with empirical observations. In practice, data scarcity or noise—common challenges in biological research—can lead to parameter sets that either fail to reproduce the desired biological dynamics (e.g., due to overfitting) or fall outside biologically plausible ranges.

In this study, we present a novel multi-objective calibration framework that integrates biological expertise to improve the optimization of dynamic models. Specifically, we employ the optimization Non-dominated Sorting Genetic Algorithm II (NSGA-II) with respect to both hard and soft constraints. Hard constraints ensure quantitative agreement with biological data, and soft constraints encode qualitative expert knowledge. These soft constraints guide the optimization by incorporating prior insights into expected system dynamics and biologically plausible parameter ranges. This approach improves the biological interpretability and reliability of the inferred parameter sets. Using a literature-based model formulated as a system of ordinary differential equations, we illustrate how the proposed framework addresses the challenges posed by high-dimensional parameter spaces and data limitations in biological modeling.

## Materials and methods

The framework consists of six main steps (Fig. 1): (1) collecting expert knowledge about the expected dynamics of the biological process under study and converting it into soft and hard constraints; (2) selecting a range for each parameter to define the search space; (3) defining a cost function to evaluate the error between biological and simulated data; (4) selecting an appropriate optimization algorithm; (5) retaining optimal solutions from the Pareto front that minimize the cost function; and (6) visualizing the dynamics of model simulations. According to the consistency of the obtained dynamics, the process is iterative and soft and hard constraints can be further modified. In summary, this strategy is designed to ensure that a biologically plausible solution is obtained by having the algorithm guided through the incorporation of all available knowledge—data, observations, and hypotheses.

**Figure 1:**
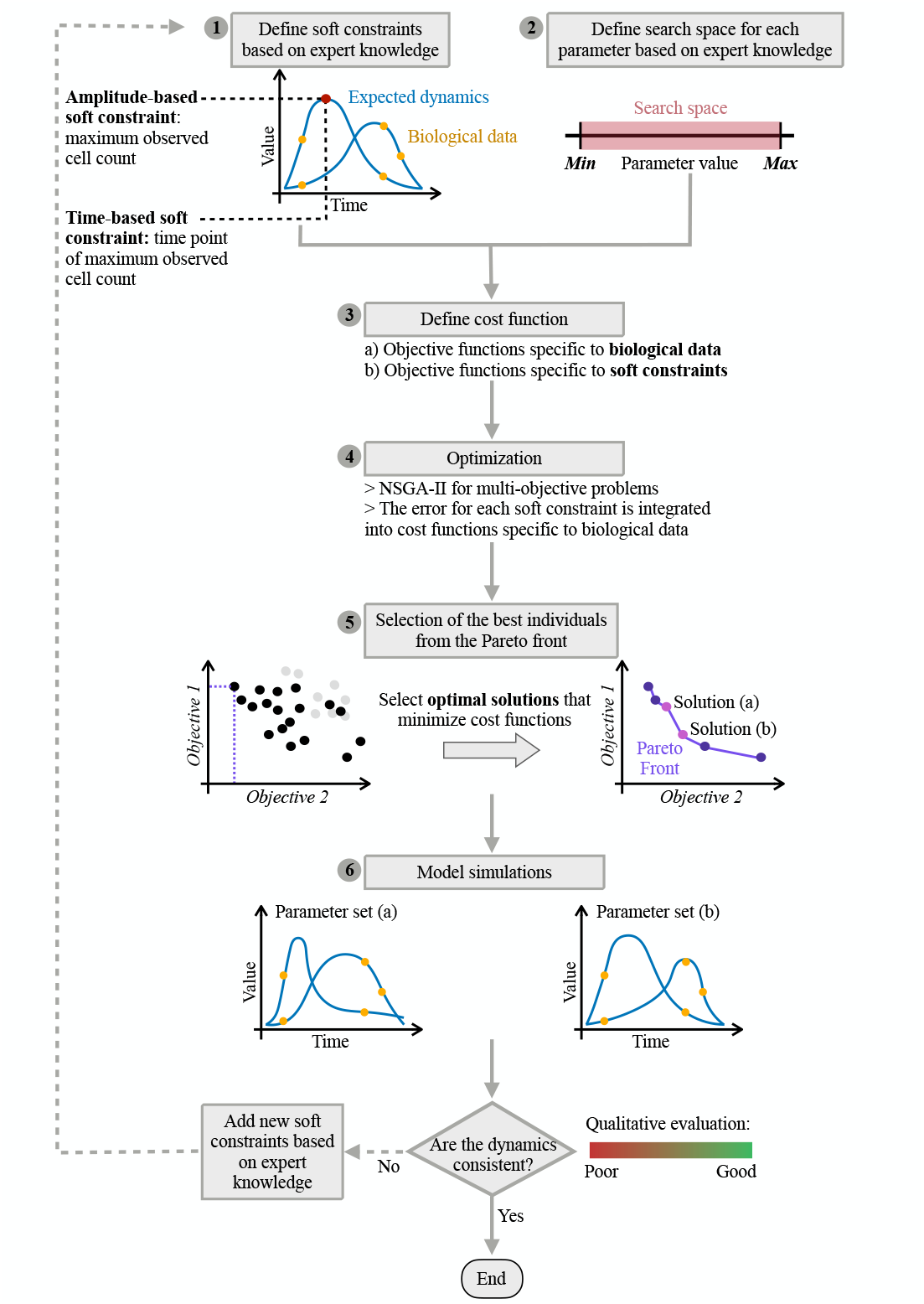
General framework of the expert-guided multi-objective optimization strategy for parameter estimation of biological systems.

### Non-dominated Sorting Genetic Algorithm optimization algorithm

NSGA-II is a state-of-the-art optimization algorithm specifically designed to handle multiple, potentially conflicting objectives [17]. For each objective to be optimized, NSGA-II requires the definition of a corresponding objective or fitness function. Unlike traditional single-objective algorithms or methods that aggregate objectives into a single scalar function (e.g., via weighted sums), NSGA-II identifies a set of optimal trade-off solutions known as the Pareto front. Each solution on this front is non-dominated, meaning that no solution in the set can be improved in one objective without incurring a loss in at least one other. This allows users to select a parameter set that best reflects their specific priorities or domain constraints.

In the context of biological modeling, NSGA-II requires fitness functions corresponding to both hard and soft constraints. Hard constraints quantify how well the model fits empirical data, typically using standard metrics such as mean squared error (MSE), mean absolute error (MAE), or similar measures. Soft constraints encode qualitative expert knowledge about the system’s expected behavior and guide the optimization process without overriding the hard constraints. Designing these soft constraints requires careful consideration to ensure they faithfully capture biologically meaningful insights before being incorporated into the optimization framework.

### Hard constraints definition

Hard constraints represent the target dynamics that the model must reproduce based on biological data specifically acquired for calibration purposes. As these data are considered ground truth, it is crucial for the model to fit the provided measurements as closely as possible. To address multi-objective optimization, we propose an alternative approach inspired from the MSE metric, defining a separate objective function for each kinetic profile. This strategy avoids the need for subjective weighting coefficients, which can be difficult to determine reliably. Furthermore, aggregating multiple kinetics into a single objective requires prior normalization to ensure that all profiles contribute comparably to the optimization process. By treating each kinetic profile as an independent objective, normalization becomes unnecessary, allowing errors to be computed and evaluated within their natural scales:

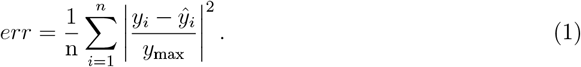

Here, *y*_*i*_ denotes the biological values, ŷ _*i*_ the simulated values, *n* the number of measurements and *y*_max_ the maximum value observed for each biological dynamic. This customized error metric ensures each error term is normalized by the maximum biological value for its respective dynamic. This normalization allows differences across profiles to be compared on a consistent scale.

### Soft constraints definition

In addition to the commonly used hard constraints reported in the literature, we propose improving the optimization process by incorporating soft constraints that encode expert knowledge about the expected system dynamics. These soft constraints may capture features such as the timing and amplitude of peak values, the sequence of events, or phase-specific characteristics of kinetic profiles. As this strategy is intended to be broadly applicable across different biological systems, the corresponding fitness functions must be carefully tailored to reflect the relevant expert insights for each specific context.

Formally, the fitness function *f*_*soft*_(*v*_*s*_) of a soft constraint can be defined as follows:

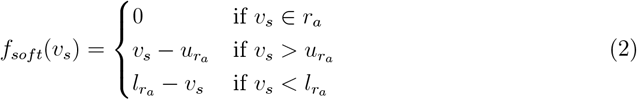

where *v*_*s*_ is the simulated data and 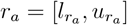 is the “acceptable range” proposed by the biological expert. The objective is to identify parameter sets that best satisfy the soft constraints, while using the hard constraints to fine-tune the parameters.

Soft constraints are often difficult to define precisely. An iterative approach is proposed, in which new constraints are progressively refined between expert-guided successive optimization loops. It is argued that the analysis of the Pareto front obtained after each loop of optimization can be used to inform the refinement of soft constraints, thereby guiding the optimization process toward increasingly biologically plausible solutions. By continuously refining or introducing additional soft constraints, this mitigates the tendency of genetic algorithms to become trapped in local optima, allowing more biologically relevant parameter sets to be identified. To avoid overly constraining the optimization process—which could reduce the algorithm’s ability to explore the parameter space—we recommend defining soft constraint functions using acceptable value ranges, considering that (i) when simulated values fall within the “acceptable range” defined in collaboration with the biological expert, the corresponding fitness value is minimal (as required by minimization algorithms); (ii) as the simulated values deviate from the nearest boundary of this range, the fitness value increases linearly.

### Define the search space for each parameter

During the optimization step, it is essential that constraints with upper and lower bounds be established for each parameter. These constraints are used to restrict the search space to reasonable ranges, thereby reducing computational time. They should ideally be defined based on available knowledge so that the algorithm is effectively guided toward biologically plausible solutions and prevented from exploring unrealistic regions.

### Case study

To evaluate the performance of the proposed strategy, we considered a benchmark model from the literature [18]. This model consists of five coupled ordinary differential equations (ODE) describing the skin wound healing process in rats over 50 days. The dynamic variables include overtime active neutrophils (*a*(*t*)), pro-inflammatory mediator (IL-6, *c*(*t*)), anti-inflammatory mediator (IL-10, *g*(*t*)), macrophages (*m*(*t*)), and apoptotic neutrophils (*n*(*t*)). The simulation was performed under the experimental condition involving 10% oil-resin. For each variable, at least one validated parameter set is available, allowing us to reproduce the original simulations and use them as a reference for evaluation (Fig. 2).

**Figure 2:**
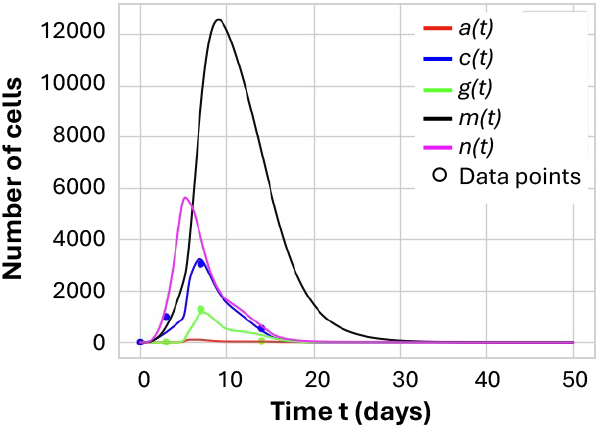
Benchmark model of skin wound healing: reproduction of simulation results from de Oliveira et al. (2021) [18]. The model simulates the kinetics of active neutrophils (*a*(*t*)), IL-6 (*c*(*t*)), IL-10 (*g*(*t*)), macrophages (*m*(*t*)) and apoptotic neutrophils (*n*(*t*)) under the experimental condition of 10% oil-resin.

The objective of this case study is to assess whether present calibration method can recover biologically plausible parameter sets capable of reproducing the benchmark simulation dynamics. Among the variables, only IL-6 (*c*(*t*)) and IL-10 (*g*(*t*)) are associated with real biological data, each comprising four time points. The optimization thus focuses on fitting these two dynamics. Thus the optimization process used the error function previously defined in Equation (1), computed independently for *c*(*t*) and *g*(*t*). The goal was to minimize the error for each kinetic profile.

Each soft constraint was implemented as an independent objective function, resulting in a multi-objective optimization problem: two for fitting biological data (*c*(*t*) and *g*(*t*), hard constraints), and several qualitative biological constraints (soft constraints). For soft constraints, only solutions that exactly satisfied the constraint (i.e., with an error equal to zero) were retained. This approach efficiently guides the optimization toward biologically valid parameter sets.

Final validation of the estimated parameters was based on the model’s ability to reproduce the full dynamics of the benchmark simulation, not just the data-associated variables. This ensures that the solution is not only data-compliant, but also biologically interpretable and dynamically consistent.

## Results

### Defining the constraints based on expert knowledge and literature

Soft constraints were derived from the dynamics described for model 1 in de Olveira et al. [18, 19], and grouped into two categories: time-based and amplitude-based constraints. Time-based constraints: (i) The peak of macrophage recruitment (*m*(*t*)) must occur between day 9 and 11; (ii) The peak of active neutrophil recruitment (*a*(*t*)) must occur before the macrophage peak; (iii) The active neutrophil peak must be transient and occur between day 3 and 8.5. Amplitude-based constraints: (i) The macrophage peak must range between 0.9 * 10^4^ and 2.0 * 10^4^ cells/*mm*^3^; (ii) The neutrophil peak must represent approximately 20–80% of the macrophage peak; (iii) The inflammatory response should be resolved by day 35, with the mean value of all dynamic variables remaining below 10 beyond this point.

These constraints are based on established biological knowledge of wound healing. The inflammatory phase involves a well-characterized sequence of immune cell recruitment, with neutrophils being the first responders, followed by macrophages [20, 21, 22]. Neutrophil recruitment is known to be driven by pro-inflammatory cytokines such as IL-6 [19], which reaches its peak around day 7 in de Oliveira’s model. Macrophage infiltration follows, becoming the dominant cell population during the transition to tissue remodeling [23, 24, 25]. The end of the inflammatory response is marked by a reduction of the number of pro-inflammatory cells, such as neutrophils, a decrease in the concentration of pro-inflammatory mediators, and a production of anti-inflammatory mediators such as IL-10. These biological insights justify the soft constraints applied in our framework. However, the experimental data on wich de Oliveira’s model is based is limited to the quantification of pro- and anti-inflammatory cytokines [19]. This may be indicative of why the soft contraintson cell type amplitude and timing do not correspond to teh literature.

### Use of the benchmark model to evaluate performance and robustness of the computational strategy

The benchmark model was used to evaluate the performance of the proposed optimization strategy, i.e. NSGA-II with 6 expert-guided constraints (NSGA-II-6C). This approach was also compared to 3 other approaches: NSGA-II with 2 expert-guided constraints (NSGA-II-2C), NSGA-II without constraints (NSGA-II-WoC), and the Genetic Algorithm (GA), commonly used for mono-objective optimization (Fig. 3). To assess the influence of the parameter search space, experiments with different levels of constraint relaxation were conducted, simulating conditions with increasing uncertainty. The initial search space was defined by applying a multiplicative factor of 100 around the nominal values reported in Oliveira et al. (e.g., for a parameter *v* = 0.3, the interval was [0.003, 30]). The parameter ranges was then progressively restricted using narrower factors of 50, 10 and 2 (Fig. 3). No prior knowledge of the true parameter values was assumed.

**Figure 3:**
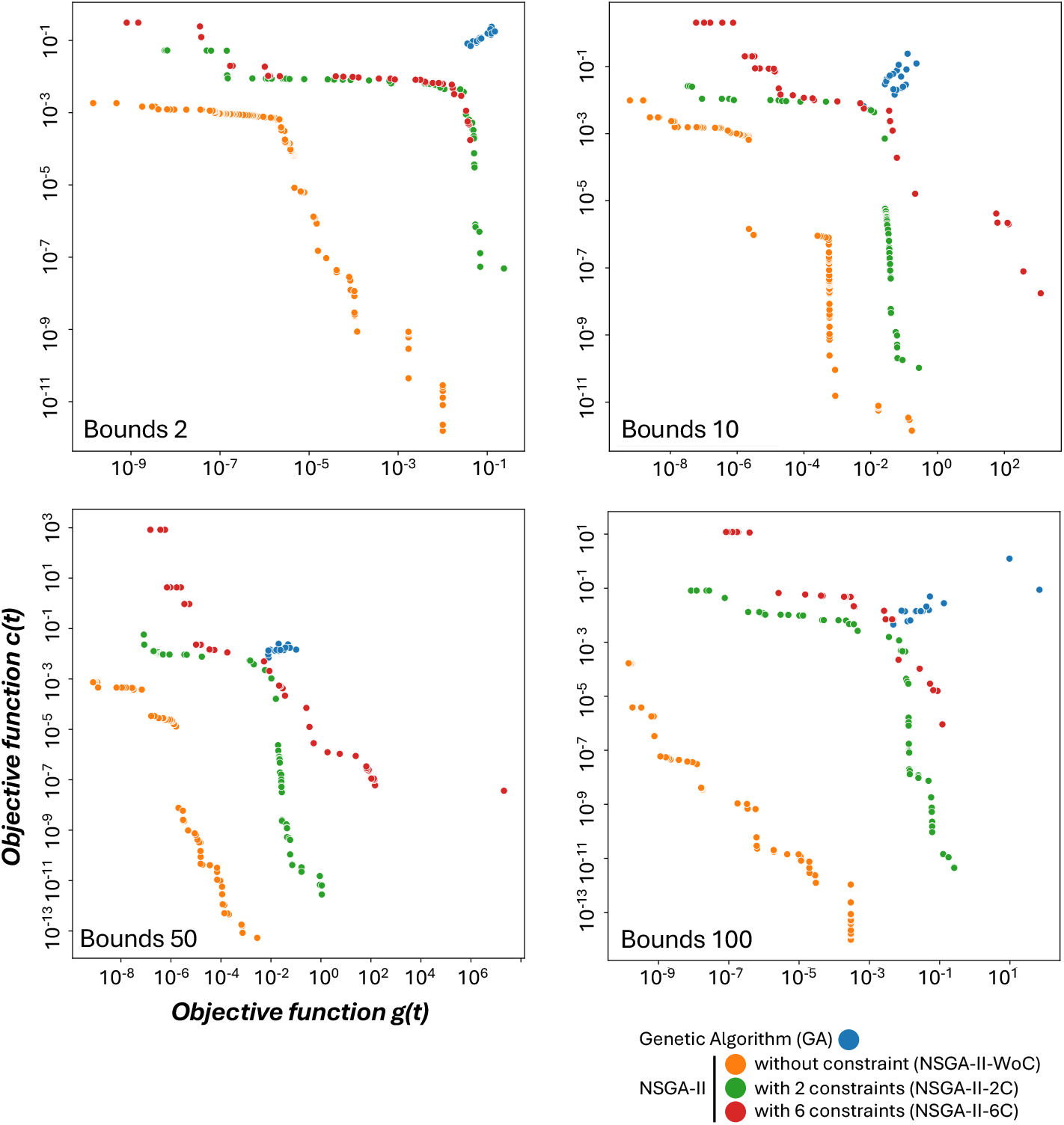
Pareto front minimizing the two objective functions *c*(*t*) and *g*(*t*). The performance of NSGA-II with 2 and 6 constraints (NSGA-II-2C and NSGA-II-6C) was compared to the optimization strategies NSGA-II without constraints (NSGA-II-WoC) and Genetic Algorithm (GA) on different search spaces. A factor 2, 10, 50 and 100 was applied to the parameter ranges.

### Adding soft constraints to optimization greatly improved the discovery of parameter sets consis- tent with biological expectations at the expense of solutions with higher objective functions

The results indicate that better trade-off solutions in terms of objective function values were identified by NSGA-II-WoC, regardless of the search space used (Fig. 3). In particular, when the search space was highly expanded (e.g., bounds scaled by a factor of 100), the best trade-off point was found using NSGA-II-WoC.

An essential step in parameter selection involves visualizing the kinetics of each variable to ensure biological consistency. To this end, the best solutions from the Pareto front of each replicate run were retained, representing the optimal trade-offs between the calibration objectives *c*(*t*) and *g*(*t*). Solutions in which the combined error for these two objectives exceeded a threshold of 1 were excluded.

When the dynamic profiles were evaluated against the expert constraints, most of the solutions were found to be inconsistent with the benchmark simulations presented in Oliveira’s work both for the GA and NSGA-II-WoC (Fig. 4). The corresponding parameter sets failed to accurately reproduce the wound healing dynamics. This discrepancy was especially evident in the expanded search space setting, where overfitting to the limited biological data (*c*(*t*) and *g*(*t*)) was likely to have occurred. In particular, oscillatory behavior was observed across both restricted and expanded search spaces (Fig. 4, Supplementary Fig. 1). Such oscillations are biologically implausible and suggest instability in the simulated system. These anomalies may result from challenges in estimating parameters in the presence of numerically stiff terms and a high-dimensional parameter space. Oscillatory dynamics suggested the presence of attractive local minima in a rugged fitness landscape. Although earlier results showed that NSGA-II-WoC yielded better Pareto front solutions in terms of objective values, the present findings confirm that the corresponding parameter sets produced dynamics inconsistent with biological expectations. This reinforces the hypothesis that overfitting to limited biological data can lead to poor generalization and failure to predict biologically realistic outcomes. The issue was exacerbated in the largest search space condition (bounds 100), which produced a significantly higher number of activated neutrophils compared to macrophages—an outcome that contradicts es- tablished biological understanding.

**Figure 4:**
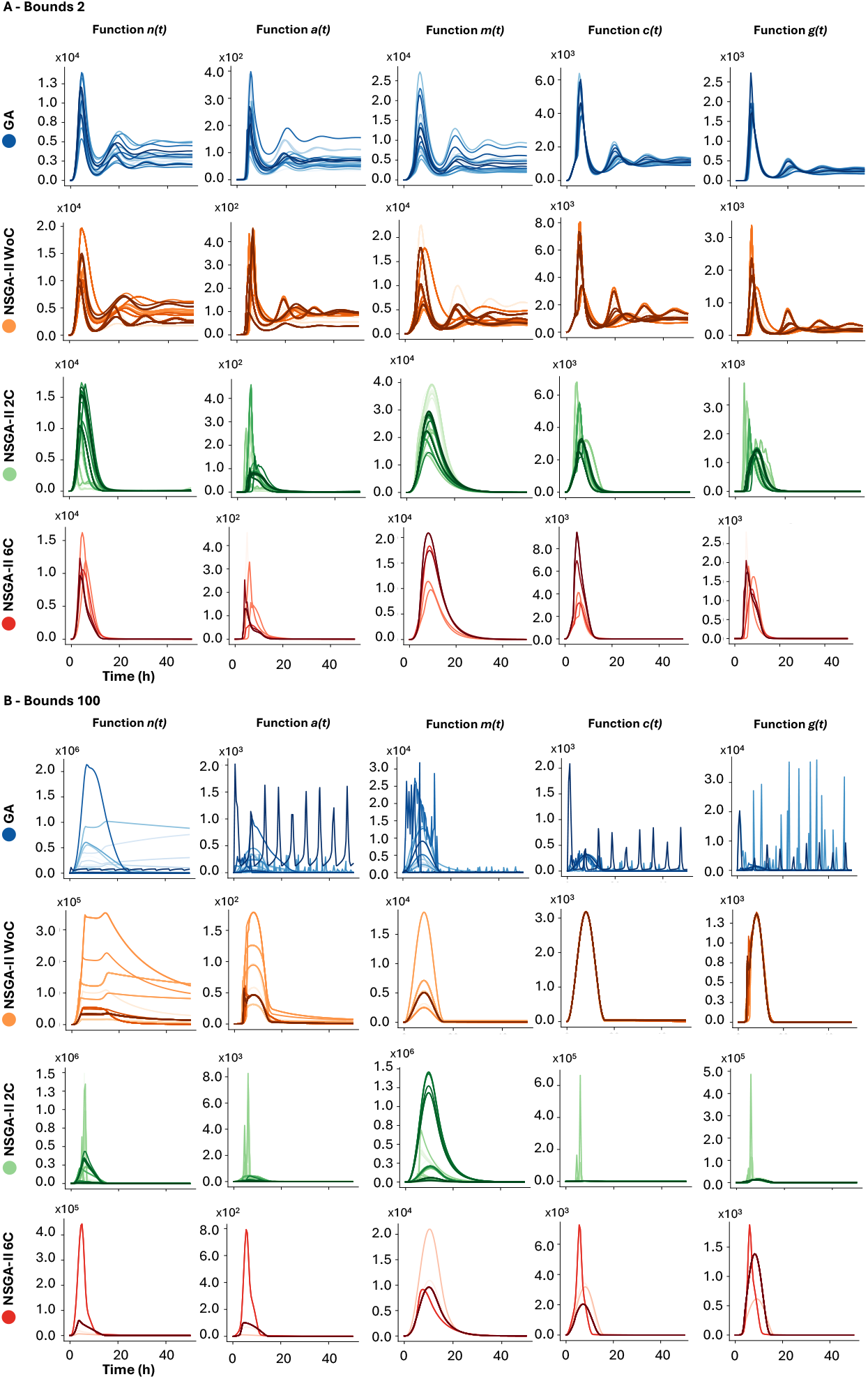
**Predicted wound healing dynamics under each optimization strategies (GA, NSGA-II- WoC, NSGA-II-2C and NSGA-II-6C). These plots show 20 replicates obtained on the pareto front, based on the minimization of the objective functions** *c*(*t*) **and** *g*(*t*). **A factor 2 (Part A) and 100 (Part B) was applied to the parameter ranges. Factors 25 and 50 can be found as Supplementary Fig. 1**.

These results underscore the limitations of optimizing solely based on sparse biological data. When expert-derived constraints were incorporated into the optimization process, the resulting dynamics were more consistent with established biological knowledge and more closely aligned with the benchmark model, regardless of the parameter space used (Fig. 4, Supplementary Fig. 1). The integration of expert guidance effectively constrained the optimization, eliminated oscillatory artifacts, and directed the algorithm toward biologically plausible solutions. These results suggest that although NSGA-II-WoC performed better in minimizing data-driven objectives, biologically implausible dynamics were often produced in the absence of expert guidance. By contrast, when expert knowledge was integrated through soft constraints, the search was more effectively directed toward parameter sets consistent with biological expectations, despite slightly higher objective values.

### Increasing the number of constraints results in improving the discovery of biologically plausible set of parameters

To further evaluate the proposed strategy, NSGA-II with constraints was tested using only two of the six previously defined expert constraints. The objective was to assess whether a biologically plausible set of parameters could still be identified with a minimal set of constraints. The selected constraints corresponded to the timing of peak recruitment for macrophages and neutrophils—temporal features that are relatively well established in the literature on wound healing. The results showed that the Pareto fronts obtained under this reduced-constraints configuration were relatively close to those obtained using the full set of six constraints, particularly for the narrower search spaces defined by bounds of 2 and 10 (Fig. 3, Fig. 4, Supplementary Fig. 1). In the case of the wider search space (bounds of 50 and 100), better objective trade-offs were observed on the Pareto front, but the resulting kinetic profiles deviated significantly from those of the benchmark model (Fig. 3, Fig. 4, Supplementary Fig. 1). While the dynamics associated with bounds of 2 and 10 were more consistent with the reference simulation, the peak amplitude of the macrophage population was substantially overestimated.

This discrepancy highlights the need for additional constraints to more accurately guide the optimization process based on domain knowledge.

Importantly, it was observed that even with only two expert constraints, solutions exhibiting the oscillatory behavior seen in the GA and NSGA-II-WoC strategies were effectively filtered out. However, two constraints alone were insufficient to consistently produce optimal solutions, particularly when the parameter search space was large. As the number of candidate parameter sets increases with broader search spaces, the likelihood of diverging from expected biological behavior also rises. In such contexts, the addition of further expert constraints becomes essential to maintain accuracy. By progressively integrating new constraints, the optimization space can be refined independently of the initial parameter bounds. This iterative strategy enables the calibration process to become increasingly focused through the incorporation of expert knowledge and qualitative judgment based on the dynamics observed. The approach can be initiated with a minimal set of constraints and extended incrementally to improve accuracy. These results demonstrate that the search space can be explored more effectively by integrating prior knowledge, observed data, or biologically grounded hypotheses directly into the optimization process.

## Discussion

Calibration is a critical step in the construction of in silico models, as it enables the reproduction of biological system behavior. However, calibrating nonlinear models remains a challenging task due to the high dimensionality of parameter spaces and the scarcity of quantitative data required for reliable parameter estimation. While quantitative measurements are typically used, qualitative knowledge and domain-specific insights are rarely leveraged. In this work, we propose an approach that integrates expert knowledge—extracted from literature, experimental observations, or mechanistic hypotheses—into the parameter estimation process of dynamic models. This knowledge is encoded as objective functions representing biological constraints that candidate parameter sets must satisfy to be considered plausible.

Biological systems are inherently complex and often admit multiple parameter sets capable of generating plausible dynamics. Inter-individual variability further contributes to this diversity. Our method embraces this multiplicity by identifying biologically coherent solutions across a wide region of the parameter space. Although a single parameter set may be selected for simplicity, it should not be considered definitive. Instead, candidate sets must be validated against independent data or through further experimental investigation. In this study, we employed NSGA-II, an efficient algorithm for bi-objective optimization. Several multi-objective optimization algorithms serve as alternatives to NSGA-II for model calibration, each with distinct strengths depending on problem complexity, dimensionality, and computational constraints. Hypothesis-based constraints could then be incorporated as independent objectives and visualized in higher-dimensional trade-off spaces. Other multiobjectives would have been possible such as NSGA-III (better suited to capture a broader and more diverse Pareto front in more than 4 objectives [26]), decomposition-based approaches such as MOEA/D (Multi-Objective Evolutionary Algorithm Based on Decomposition wich divides problems into scalar subproblems for parallel optimization [27]), GDE3 (Generalized Differential Evolution 3, combining differential evolution with Pareto dominance [28]), hybrid stochastic-gradient methods such as spMODE-II (integrating differential evolution with sensitivity analysis [29]), particle swarm and metaheuristic variants such as MOPSO (Multi-Objective Particle Swarm Optimization, using particle swarms for exploration [30]). NSGA-II stands out as a versatile, efficient, and user-friendly algorithm for multi-objective calibration of mechanistic models [31, 32]. Its ability to generate diverse Pareto-optimal solutions supports a nuanced understanding of parameter uncertainty and model trade-offs. While alternatives may excel in specific contexts, NSGA-II’s balance of performance, ease of use, and robustness makes it a preferred choice in many practical calibration scenarios. Incorporating NSGA-II with complementary uncertainty quantification techniques can further enhance confidence in calibrated models, to avoid that the exploration of solutions will be restricted to local minima. For example, it is possible to refine the Pareto front using posterior distributions from Bayesian inference [33], modifiy the crowding distance to prioritize parameters in sparsely populated regions of the Pareto front [34] or improve the diversity of the initial generated population [35]. Recognizing the computational limits of NSGA-II handling more than 3 objectives, a meta-objective approach may enhances NSGA-II’s ability to handle many-objective optimization problems (four or more objectives) by reformulating the problem into a new “meta-objective” space [36].

The integration of expert-guided qualitative constraints for model calibration offers substantial benefits that enhance the calibration process beyond purely quantitative optimization [37, 38]. Expert knowledge provides qualitative constraints or constraints-such as expected behavioral patterns, or known system dynamics-that help steer the NSGA-II search toward more meaningful regions of the parameter space. This guidance reduces the risk of converging to parameter sets that fit data numerically but lack biological realism [39, 40]. This multi- constraints approach also facilitates the detection of prominent behavioral solutions that satisfy both quantitative and qualitative expectations, leading to more robust and interpretable calibrated models, narrow the Pareto front to solutions that are both statistically sound and scientifically plausible thus improving model generalizability, reduce uncertainty and facilitates interpretability [39, 40]. The proposed strategy is both flexible and iterative, enables the integration of any type of knowledge-based criteria. Constraints can be ranked by confidence levels based on biological evidence, and progressively introduced to refine the calibration process. For example, we initially used only two temporal constraints—based on the well-characterized dynamics of macrophages and neutrophils—yet this minimal set was insufficient to guide the algorithm toward the target solution. Adding further qualitative biological constraints acts as a form of regularization, limiting overfitting. It enables the generation of biologically plausible dynamics even under conditions of high uncertainty, particularly when the parameter space is extremely broad (e.g., 100-fold). This supports the idea that the framework can be generalized to other systems with limited experimental information.

While this strategy enables the generation of calibrated models that capture key biological dynamics, model validation and structural identifiability remain crucial. Further anal- yses—such as sensitivity analysis—are required to assess parameter uncertainty, particularly when dealing with limited data and high-dimensional models [41, 42]. These analyses may also inform the design of new biological experiments aimed at constraining the most sensitive parameters. To effectively explore the range of solutions that a model can produce, flexible constraint definitions are essential. The parameters obtained should be further validated *in vitro* with new biological data.

This work introduces a methodological contribution that lies in the hybridization of a multi-objective optimization algorithm with iteratively integrated expert knowledge. This hybrid framework goes beyond the limitations of purely statistical or Bayesian approaches, which often struggle to capture biologically realistic behavior in complex, underdetermined systems. The strategy has been validated on a published benchmark model, demonstrating its applicability and robustness. Notably, our results show that even a small number of well-chosen qualitative constraints can substantially improve simulation outcomes, although they remain insufficient on their own. This observation highlights the need to address the trade-off between selectivity and generality of constraints. It also opens the door to new research directions, such as identifying the most informative constraints, designing constraint profiles tailored to specific biological processes, and formalizing these elements as part of a modular, reusable library. Building on this idea, the framework could be extended into a semi-supervised iterative process, in which constraints are progressively refined either manually by experts or automatically through learning from a set of labeled, biologically plausible simulations. Ultimately, this approach reframes modeling as a knowledge-driven, exploratory endeavor. Its general methodology can be adapted to a variety of modeling paradigms, including agent-based models.

Despite its promising results, the present approach has certain limitations that warrant discussion. The benchmark model used for validation is relatively simple, consisting of five coupled ordinary differential equations with no delays or feedback loops. It remains to be seen how well the strategy scales to more complex systems involving dozens of equations, intricate regulatory interactions, or time delays—features commonly encountered in fields such as immunology, pharmacokinetics, or tumor growth modeling. Additionally, the expert constraints employed in this study were defined heuristically, based on literature knowledge and biological intuition. A more systematic formalization—through probabilistic encoding or expert-driven annotation—could enhance the reproducibility and robustness of the approach. Future extensions could include the integration of automatic validation tools, such as dynamic pattern classifiers [43], to objectively evaluate the plausibility of simulated trajectories.

Building upon the proposed framework, several avenues can be explored to further improve its applicability and generalizability. First, developing a modular library of expert criteria tailored to different biological processes—such as immunology, wound healing, or oncology—would facilitate reuse and standardization across modeling efforts. In parallel, automated strategies for identifying the most informative constraints could enhance calibration efficiency and reduce expert burden. Designing constraint profiles specific to each application domain will also improve biological relevance and interpretability. A natural extension of this work involves framing the calibration process as a semi-supervised iterative loop, combining expert interventions with automated learning from biologically validated simulations. Integrating dynamic pattern classifiers or other automated validation tools would allow for objective assessment of the plausibility of simulated trajectories. Moreover, the proposed framework could serve as a pre-processing step for more computationally intensive Bayesian inference methods, by narrowing the parameter space to biologically relevant regions. Ultimately, translating this approach into a dedicated software platform—combining NSGA-II, interactive trajectory visualization, and expert-guided constraint integration—would broaden its accessibility. The methodology is also compatible with other modeling paradigms, such as agent-based models, where qualitative dynamics are often essential yet difficult to quantify, making expert-informed calibration particularly valuable.

## Conclusion

We introduced an expert-guided multi-objective optimization framework that combines quantitative data with qualitative biological knowledge to calibrate mechanistic models. Using NSGA-II, our approach encodes expert constraints as soft constraints, improving convergence towards biologically plausible parameter sets, even under data scarcity or wide search spaces. Comparative evaluations showed that unconstrained optimization, while achieving lower numerical errors, often produced biologically implausible dynamics. In contrast, expert-informed calibration acted as an effective regularizer, filtering out unrealistic behaviors. The iterative integration of qualitative constraints allowed for progressive refinement of the search space, offering a practical strategy to guide the optimization towards coherent solutions. While demonstrated on a moderately complex benchmark, the framework is extensible to more intricate models. Future work will focus on scaling the method, formalizing expert input, and integrating it into semi-automated calibration pipelines. By leveraging expert knowledge in addition to sparse data, this approach supports the development of reliable models in contexts where data alone is technically complex to acquire.

## Aknowledgment

This work was supported by the Occitanie Region, the Federal University of Toulouse Midi-Pyrénées (grant ADI, N°ALDOCT-000689/19008724). This study has been partially supported by the Agence Nationale pour la Recherche, through the grant EUR CARe N°ANR-18-EURE- 0003, the grant N°ANR-ANR-24-CE45-5810 and the national infrastructure “ECELLFrance: Development of mesenchymal stem cell based therapies” (PIA-11-INBS-0005) in the framework of the Programme des Investissements d’Avenir.

## Code availability

https://gitlab.com/isicell-irit/expert-guided-multi-objective-optimization

## Authors’ contributions

L.DCF., D.B., S.CB., P.M. and B.C. conceived the experiments. L.DCF. and D.B. conducted the experiments. L.DCF., D.B., S.CB. and P.M. analyzed the results. S.CB., P.M. and B.C. wrote the manuscrit. L.DCF., D.B., S.CB., F.P., P.M. and B.C. reviewed the manuscrit.

## Footnotes

1 These authors contributed equally to this work

2 These authors contributed equally to this work

