## Supplementary figures and images for "Expert-guided multi-objective optimization: an efficient strategy for parameter estimation of biological systems with limited data"

### Supplementary Fig. 1

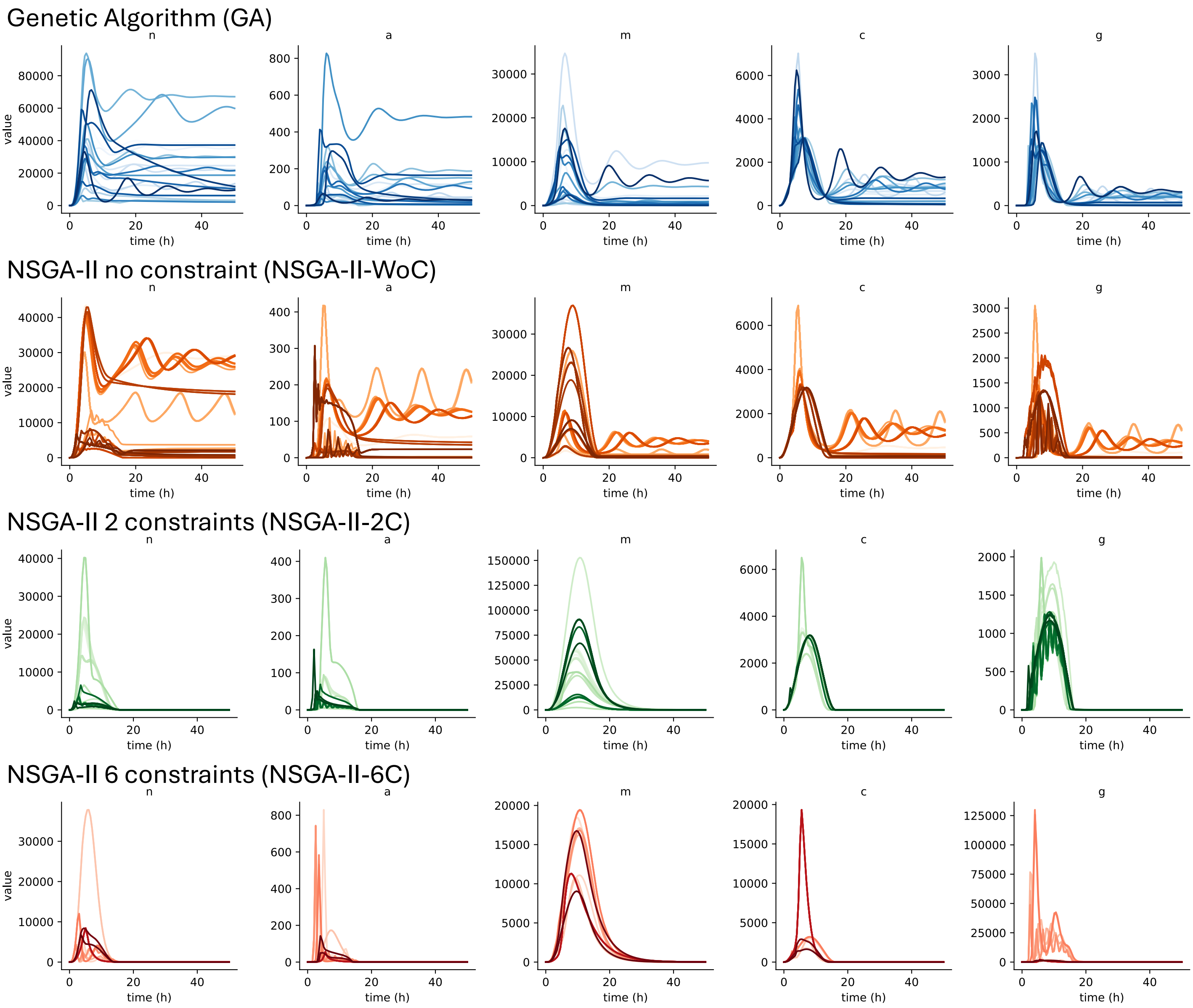

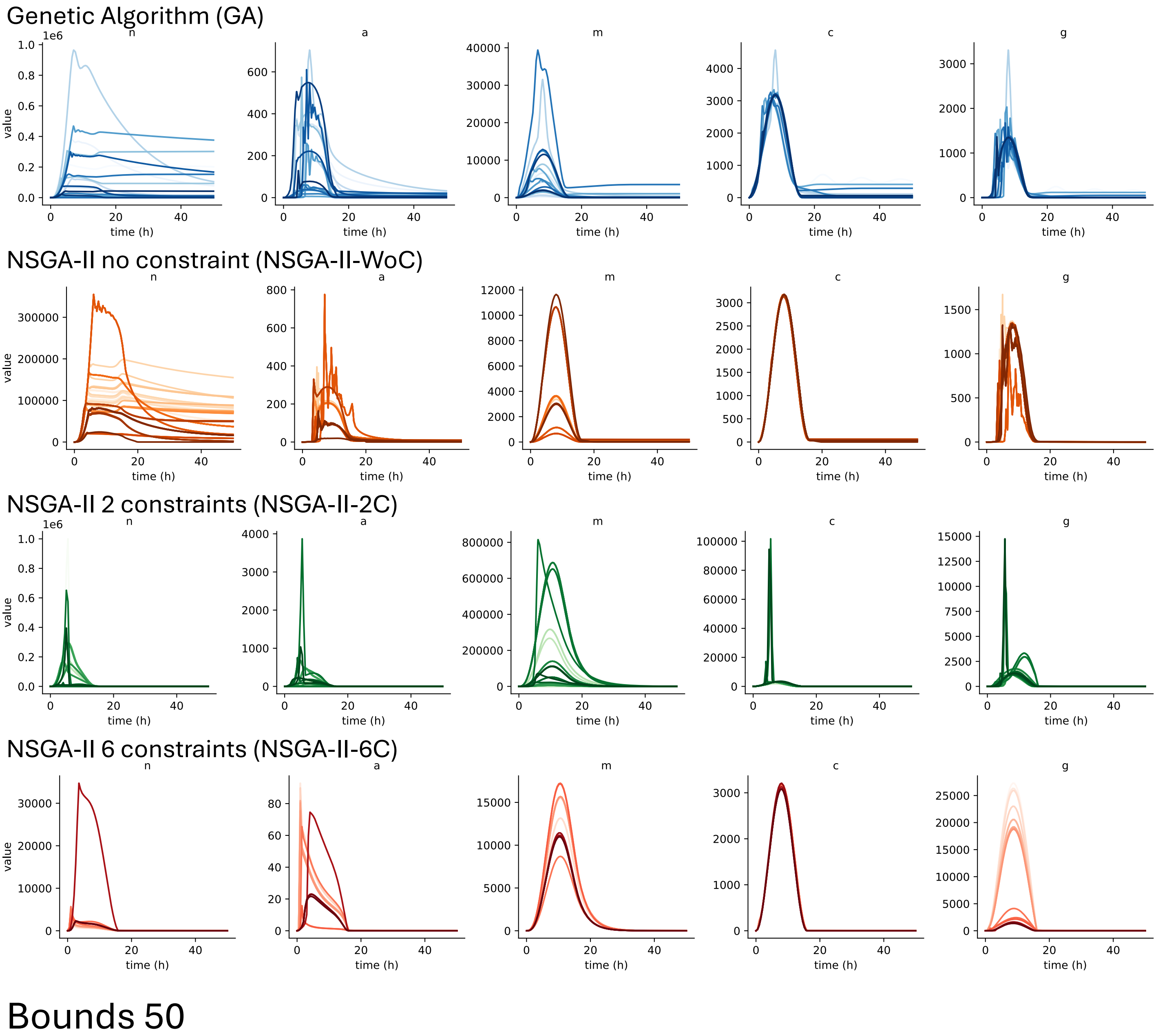
